# Metabolomic differentiation of amino acid profiles in adult deltamethrin-resistant *Aedes albopictus* (L.)

**DOI:** 10.1101/2024.07.19.604257

**Authors:** Song-Quan Ong, Intan H Ishak, Gomesh Nair, Rolinus Paulous

## Abstract

Understanding the susceptibility status of mosquitoes to insecticides is critical for effective decision making regarding the use or rotation of insecticides in control programs. In this study, we demonstrated the use of amino acid profiling for the detection of deltamethrin-resistant *Aedes albopictus* (L.). Mosquitoes collected in the field were first tested with WHO adulticide bioassay kits, and the amino acid profiles of the resistant mosquitoes were compared with the susceptible strain of *Ae. albopictus*. Samples were lyophilized and derived by silylation and then analyzed by gas chromatography-mass spectrometry (GC-MS). Using standardized, known concentrations of amino acids, we quantified the amino acids in both resistant and susceptible strains. An independent t-test was performed to compare the concentrations of each amino acid between strains. Logistic regression was then performed to assess the relationship between amino acid concentrations and susceptibility status of the mosquitoes. Our results showed that the amino acids in resistant mosquitoes differed significantly from those in susceptible mosquitoes, with the exception of serine. Further regression analysis showed that seven amino acids significantly predicted susceptibility, suggesting that they are suitable as biological indicators for rapid assessment of resistance status in field mosquitoes.

**Graphic abstract:** 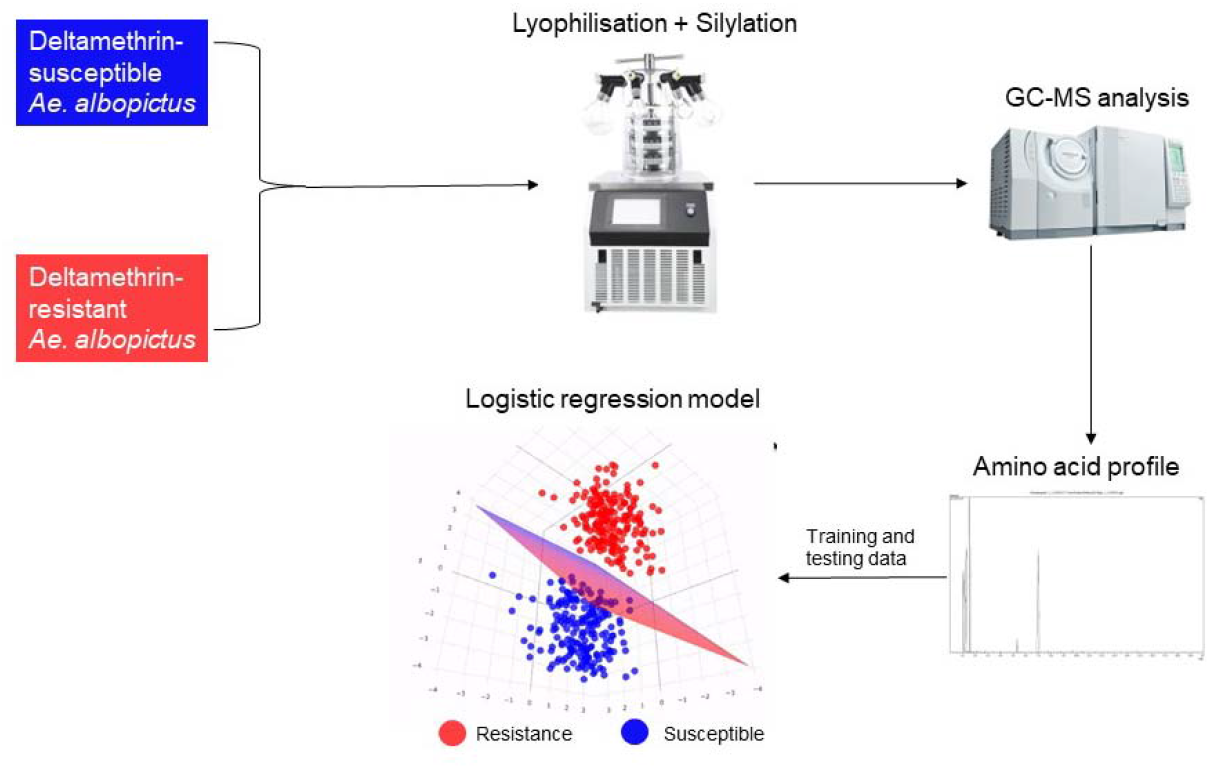

Identification of deltamethrin-resistant mosquitoes based on differences in the amino acid profile: Deltamethrin-susceptible and -resistant mosquito strains were lyophilised and converted into esters by silylation, which were then analysed using a gas chromatography-mass spectrometer (GC-MS). The difference between susceptible and resistant could be classified by developing a classification model with logistic regression.

## Introduction

The dengue virus, which is mainly transmitted by *Aedes* mosquitoes, is a major health problem worldwide and causes a spectrum of diseases ranging from mild fever to life-threatening conditions (Ross, 2010). The virus has four serotypes and has evolved rapidly as it has spread around the world (Ross, 2010). Factors influencing dengue virus transmission include geography, climate, urbanization and globalization (Hoque, 2023). Each year, 50-100 million new infections occur, resulting in approximately 20,000 deaths (Islam et al., 2021). Prevention and control strategies focus on vector management as there is no effective tetravalent vaccine or specific treatment (Sarwar, 2014). Integrated approaches to mosquito control include household and large-scale interventions, sustainable control measures and coordinated multi-sectoral interventions such as early detection, appropriate medical care and increased public awareness can help reduce mortality rates (Ong, 2016, Ong and Hamdan 2024). Nevertheless, the challenges of mosquito control remain and resistance to insecticides could likely be one of the major drawbacks of control programs (Coleman & Hemingway, 2007, Karunaratne et al., 2018).

Insecticide resistance is a major threat to mosquito control programs as there are currently no suitable substitutes that could be used to rapidly control mosquito populations (Liu 2015). This is particularly true for pyrethroids such as deltamethrin, which are often used as residual sprays in urban areas to control dengue vectors — *Aedes aegypti* (L.) and *Aedes albopictus* (L.) (Smith et al 2016). Deltamethrin is one of the six pyrethroid insecticides recommended by the World Health Organization (WHO) for the treatment of mosquito nets as part of the WHO evaluation scheme for pesticides (WHO, 2017).

Several studies have reported the development of resistance to deltamethrin in these mosquitoes due to the involvement of monooxygenases, hydrolytic esterase or a knockdown resistance (kdr)-based mechanism (Ranson et al., 2011). Zulfa et al. (2022) reported that the prevalence of deltamethrin resistance in *Ae. albopictus* was 2% between 2011 and 2021, which is relatively low compared to malathion (21%), DDT (64%) and permethrin (6%). However, the level of resistance is expected to increase as deltamethrin is used more frequently in mosquito populations.

Although efforts are being made to develop new insecticides that effectively control insect strains resistant to currently used insecticides, the need to protect and extend the useful life of current insecticides remains critical (Sparks and Nauen 2015). For this reason, resistance management must be given a higher priority in decision-making in mosquito control programs than is currently the case. Currently, the insecticide resistance surveillance program relies on the discriminatory concentrations provided by the WHO (WHO, 2023). According to this, a mortality rate of 98-100% indicates susceptibility, while a mortality rate of 80-97% indicates possible resistance that needs to be confirmed. However, this method is often time-consuming and labor-intensive, as field strains must be bred in sufficient quantities and tested with WHO test kits.

The search for alternatives to detect insecticide resistance has led to research into various biochemical and molecular markers. Among these, metabolite analysis by amino acid profiling has emerged as a suitable approach as it is able to reflect metabolic changes associated with resistance mechanisms. Insecticide resistance is often associated with complex biochemical processes that alter the normal physiology of mosquitoes. One of the most important mechanisms is metabolic detoxification, in which specific enzymes break down insecticides before they can exert their toxic effects. These enzymes include esterase, glutathione S-transferases (GSTs) and cytochrome P450 monooxygenases (P450s), which are involved in the detoxification process (Bosch-Serra et al 2021). The production and activity of these enzymes is closely linked to the metabolic pathways in which amino acids are involved. Amino acids serve as precursors for the synthesis of detoxification enzymes. For example, glutathione (one of the members of the GST), an important molecule in the detoxification process, is synthesized from amino acids such as glutamate, cysteine and glycine (Forman et al. 2009). Changes in the concentrations of these amino acids may indicate an upregulation of detoxification pathways, suggesting insecticide resistance. Studies have shown that resistant mosquitoes often have altered amino acid profiles that can be detected and used as markers of resistance (Beech et al. 2011, Martin-Park et al. 2017). Several studies have shown the potential of amino acid profiles for the detection of insecticide resistance. For example, Martin-Park et al. (2017) analyzed the amino acid composition of *Culex quinquefasciatus* and found significant differences between susceptible and resistant strains when exposed to different insecticides. Similarly, Huang et al. (2020) identified 17 amino acids associated with deltamethrin resistance in *Ae. albopictus*, highlighting specific amino acids that were significantly elevated in resistant strains.

In addition, advances in analytical chemistry and metabolomics have provided powerful tools for amino acid profiling. Techniques such as gas chromatography-mass spectrometry (GC-MS) and liquid chromatography-mass spectrometry (LC-MS) allow the precise quantification and identification of amino acids in mosquito samples (Kvitvang et al. 2011). These techniques have been successfully used in various studies to differentiate between resistant and susceptible mosquito populations based on their metabolic profiles (Faucon et al., 2017). Therefore, in this study, we aim to utilize the amino acid profiles to develop a diagnostic tool for deltamethrin resistance in *Ae. albopictus*. By comparing the amino acid profiles of resistant and susceptible strains, we aim to identify important amino acids that can serve as biomarkers for resistance.

## Materials and methods

### Chemicals

All chemicals used in this study were sourced from Sigma-Aldrich Malaysia. Specifically, the amino acid standards (Supelco, AAS18 analytical standard); pyridine (Sigma-Aldrich, Ph. Eur > 99.5% for HPLC and GC). 2-Methyl-1-propanol (Sigma-Aldrich, 99.5%, for GC), chloroform (Sigma-Aldrich, >99.9%, for HPLC and GC) and acetonitrile (Sigma-Aldrich, >99.9%, for HPLC), HCl (Merck, 1M and 6M in H2O) and N-methyl-N-(tert butyldimethylsilyl) trifluoroacetamide (MTBSTFA) were purchased from Sigma-Aldrich (Supelco, 98.5%).

### Mosquitoes

Adult female mosquitoes were collected from March to April 2022 from three locations in Penang, Malaysia – Balik Pulau (5.352000771350706, 100.23580890621383), Glugor (5.348699234705175, 100.29572394873853) and Sungai Ara (5.334935076839994, 100.28022835371894) for insecticide trials. The choice of location was based on the area of the dengue hotspots and the usual treatment with insecticides there. The mosquitoes were caught using the Bare Leg Catch (BLC) and were reared in another generation to reach the number of adult mosquitoes for the WHO insecticide susceptibility bioassays. The mosquito was reared at the Vector Control Research Unit (VCRU) of Universiti Sains Malaysia. The larvae are reared in dechlorinated water and fed with laboratory diet (dog biscuits: yeast: milk powder: bovine liver powder in 3:1:1:1 ratio). The pupae are placed in a 30 × 30 × 30 cm net cage from which the adults hatch. The adult mosquitoes are fed 10% sucrose mixed with a vitamin B complex as an energy source. For the bioassay, three-to-five-day old adult female mosquitoes were used that were not fed with blood.

### WHO insecticide susceptibility bioassays

Using WHO (2016, 2023) recommended procedures, a complete assay included four replicates with 20 field-caught adults per tube and two control replicates with 20 susceptible *Aedes albopictus* mosquitoes per tube. A total of 200 female mosquitoes were exposed to the WHO bioassays, while 200 mosquitoes were used as controls. Adult female mosquitoes were exposed to insecticidal papers impregnated with deltamethrin (0.03%) in WHO tubes for one hour (WHO 2016). Two control tubes, each lined with paper impregnated with silicone oil, were carried out at the same time as the insecticide tests. After exposure, the mosquitoes were treated with a 10% sugar solution and kept at 24±2 °C and 73±3% humidity. The mortality of the mosquitoes was observed and recorded after 24 hours. The observed mortality of the test sample was calculated by summing the number of dead mosquitoes across all exposure tubes and then expressed as a percentage of the total number of exposed mosquitoes. Mosquitoes were stored at -40°C for analysis of animo acids

### Metabolomics analysis

The amino acid profile was slightly modified following Sobolevsky et al. (2003) and Jiménez-Martín et al. (2012). Standard solutions of the amino acids used for silylation were prepared in 10 mL of 0.1 and 0.01 M HCl at a concentration of 10 – 7 g/µL each. 100 µL of each solution was evaporated to dryness at room temperature in a gentle stream of nitrogen. The remaining water was removed with methylene chloride. Derivatisation of the dry residue was carried out with MTBSTFA. 100 µL of acetonitrile and 100 µL of MTBSTFA were added to the residue. After sonication for 30 seconds, the mixture was heated to 70°C for 30 minutes. 1 µL of the reaction mixture was injected into the GC-MS.

The mosquito samples were pooled into at least four samples per tube and freeze-dried. The freeze-dried samples were treated with 500 µL of double distilled water added to 100 mg of the lyophilizate. The mixture was sonicated for 5 minutes and then centrifuged at 7000 rpm. 500 µL of the yellowish transparent solution above the undissolved residue was transferred to a vial containing 150 µL of isobutanol, 50 µL of pyridine and 150 µL of isobutylchloroformic acid were added and the mixture was sonicated for 30 s. The derivatives formed were then treated with double distilled water. The derivatives formed were then extracted with 500 µL of chloroform by vigorous shaking and subsequent centrifugation of the solution. 1 µL of the chloroform layer was injected into the GC-MS.

GC-MS analysis of the amino acid was performed using a GC-MS (Shimadzu QP-2010 gas chromatograph attached to a Shimadzu GCMS QP-2010 plus detector (Shimadzu Corp., Japan)) with the GC column SGE BPX-5 (30.0 m X 0.25 μm i.d., layer thickness 0.25 μm) according to the method described by Nyberg (1986) with some modifications. For GCMS detection, an electron ionization system with ionization energy of 70 eV of was operated in EI mode and an acquisition mass range from 50 to 550 Da at 0.25 scan s-1. A 1 μL sample of chloroform layer was injected in split mode with a split ratio of 1:50 using a Shimadzu AOC-5000 auto-injector. Specific experimental parameters included a helium flow rate of 1 mL/min, an injector temperature of 250 °C, an initial GC temperature of 140 °C with a hold time of 2 minutes, and a GC ramp from 8 °C/min to 300 °C with a hold time of 5 minutes. The ion source and interphase of MS were set at 200 °C and 250 °C, respectively.

The detection database was based on the National Institute of Standards and Technology (NIST) Mass Spectrometry Data Centre (NIST 2017) and the concentration of amino acids in ppm was calculated based on the peak area of the standard and the sample. The retention indices were calibrated using a homologous series of n-alkanes (C8 - C40) (Custom Retention Time Index Standard, Restek Corp, United States) as an external standard to determine retention indices under same parameters prior to sample analysis.

### Statistical analysis

A Shapiro-Wilk normality test was performed for each amino acid to assess the nature of the data. To compare amino acid levels between resistant and susceptible strains of *Aedes albopictus*, the Mann-Whitney U test with a significance level of p = 0.05 was used. A binomial logistic regression model with a split of 70% train data and 30% test data was also created to analyse the relationship between the amino acid concentration and the resistance status of the mosquitoes, using a significance level of p = 0.05.

## Result

A total of 1600 mosquitoes from two different generations were used for the WHO bioassay tests. The resistance status of 440 mosquitoes was confirmed and extractions and GC-MS analyses were performed. A total of twelve amino acids were detected in both the susceptible and resistant mosquito strains. The Shapiro-Wilk normality result with a p-value of 0.66 indicates the non-parametric nature of the data. Therefore, a Mann-Whitney U-test with a significance level of p = 0.05 was used to compare the amino acid concentration between susceptible and resistant strains. Table 1 shows the amino acid profile and the comparison between the two strains. In general, serine is the only amino acid that is not significantly different between susceptible and resistant strains; eight amino acids were significantly higher in the susceptible strains than in the resistant strains. Of note, three amino acids (Ala, Val and Gly) were significantly higher in the resistant strains than in the susceptible strains, indicating their role in the assembly of the detoxification enzyme.

**Table 1.**
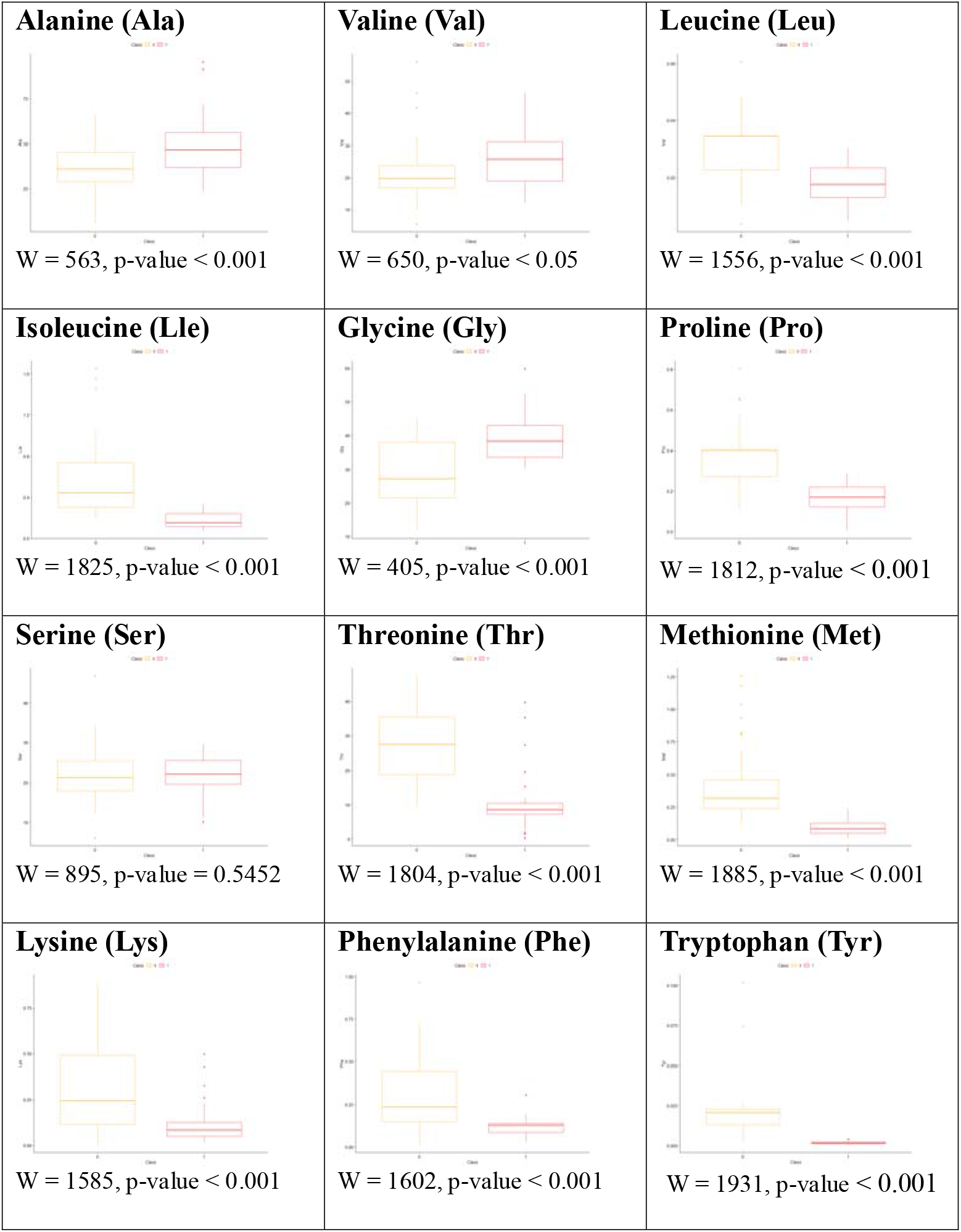
Boxplot for twelve amino acids, where class “0” stands for susceptible and class “1” for resistant, with n = 88 Wilcoxon (w) and p-value; the x-axis stands for the classes, the y-axis for the concentration of the amino acids in parts per million (ppm)

In logistic regression, a low value of the Akaike information criterion (AIC) – 26 - indicates a high probability that the model fits the training data set. The logistic regression coefficients and likelihood show that seven amino acids were significantly associated with the prediction of mosquito resistance status, with Ala, Val, Met and Tyr at p<0.05 and Pro, Cly and Thr at p<0.001 (Table 2).

**Table 2.** Logistic regression analysis of amino acids concentration to predict the resistance status of *Aedes albopictus*.

| <u>Amino acid</u> | <u>Coefficient (b)</u> | <u>se</u> | <u>z</u> | <u>Prob.</u> |
| --- | --- | --- | --- | --- |
| Ala | 0.004 | 0.002 | 2.143 | .035* |
| Val | -0.006 | 0.003 | 2.209 | .030* |
| Leu | -0.755 | 2.858 | 0.264 | .793 |
| Pro | -1.038 | 0.217 | 4.791 | .000** |
| Lle | -0.087 | 0.218 | 0.400 | .691 |
| Gly | 0.011 | 0.003 | 3.501 | .001** |
| Ser | 0.004 | 0.004 | 0.897 | .373 |
| Thr | -0.011 | 0.003 | 3.858 | .000** |
| Met | -0.472 | 0.218 | 2.164 | .034* |
| Phe | 0.170 | 0.261 | 0.652 | .516 |
| Lys | -0.321 | 0.175 | 1.837 | .070 |
| Tyr | -5.437 | 2.177 | 2.497 | .015* |

b- logistic regression coefficient; se- standard error; z- z-ratio; prob- probability; AIC = 26, Null deviance: 1.22 x 102, df = 87; \*significance level $p < 0.05$ ; \*\*significance level $p < 0.001$

## Discussion

The results of this study emphasise the metabolic differences between deltamethrin-resistant and susceptible strains of *Aedes albopictus* mosquitoes. The resistant strain had a different amino acid composition, which is consistent with the theory that these mosquitoes adapt their metabolism to be resistant to insecticides. In particular, the resistant mosquitoes had significantly higher levels of three amino acids (Ala, Val and Gly) compared to the susceptible strain. These amino acids are likely involved in the synthesis of detoxification enzymes such as glutathione S-transferase (GST), which play a crucial role in the neutralization of insecticides.

Our findings that the resistant strain of *Ae. albopictus* has a high concentration of glycine are consistent with previous studies by Hayes et al. 2005 and Liu et al. (2021). They investigated the metabolic pathway of GSTs in relation to toxic substances, including insecticides, and found that the GSTs catalyse the conjugation of reduced glutathione (GSH), which is a nucleophilic tripeptide containing glycine as one of the amino acids. Consequently, the higher activity of GST also leads to a higher concentration of glycine. A similar result to ours, that the serine concentration is not significant between the susceptible and resistant strain, was also confirmed by Adedeji et al. 2020. They investigated the significance of the use of metabolic proteins as insecticides and examined the arrangement of the amino acid sequences of AChE from 13 animal species, including *Anopheles gambiae, Culex pipiens* and *Aedes aegypti*, and found that serine is generally an animo acid residue that is conserved in most animals. This suggests that serine is highly involved in vital metabolic and physiological processes such as blood digestion. Our result extends the study of Huang, in which the deltamethrin-resistant strain of *Ae. albopictus* larvae were analyzed, which differs significantly from the susceptible strain in nine animo acids, and our result additionally includes one animo acid (methionine) that could serve as a biomarker to determine resistance status. Furthermore, our study fills the gap of Huang et al. (2020), in which only mosquito larvae were analyzed, while the adult sample we used was more frequently exposed to deltamethrin.

Interestingly, our study also identified eight amino acids that were significantly lower in the resistant strain compared to the susceptible strain. This discrepancy could be attributed to the genetic and metagenomic adaptations of the resistant mosquitoes, which prioritize the production of detoxification enzymes over other metabolic functions. To our knowledge, this is the first study to report on the amino acid profile of deltamethrin-resistant adult female *Ae. albopictus*, providing a new perspective on resistance mechanisms. Deltamethrin was selected for this study because it is commonly used in mosquito control programs. An innovative approach we used to discriminate between susceptible and resistant strains was logistic regression, a machine learning technique. This method, which has not previously been used in metabolomics studies of insecticide resistance, proved to be effective in classifying resistance levels based on amino acid profiles. Our model showed high predictive accuracy, as indicated by a low Akaike Information Criterion (AIC) score of 26.

This approach offers three major advantages over conventional dose-response bioassays. Firstly, the higher sensitivity and specificity that we owe to the amino acid profiling technique, that can detect down to the ppm range and is able to detect subtle changes in metabolic pathways, providing precise markers of resistance (Rampler et al., 2020). Secondly, it is the fast and high throughput that the demonstrated method allows for the efficient screening of large numbers of samples (Dueñas et al., 2023), covering a wider area of investigation. Thirdly, workplace safety, as the technician is only exposed to a minimal amount of insecticide and the complexity and labour intensity of resistance monitoring is reduced (Damalas & Eleftherohorinos 2011).

Based on our logistic regression result, it is possible to identify specific amino acids associated with resistance and allow prediction of resistance development before it reaches a critical level so that timely intervention can be made (Ding et al., 2023). However, this study has several limitations. Firstly, there are spatial and temporal variations, as the study used a mosquito population from a specific geographical area and at a specific time point, which may not represent the genetic diversity of *Ae. albopictus* worldwide. Secondly, the complexity of the resistance mechanisms must be taken into account. Amino acid profiling may not capture all resistance mechanisms, especially other resistance mechanisms such as target site insensitivity or cuticle thickening. The integration of other “omics” approaches, such as genomics and proteomics, may provide a more comprehensive understanding of resistance. By recognising these limitations, future research can build on current findings to develop more robust and comprehensive strategies for the surveillance and management of insecticide resistance in mosquito populations.

## Ethical declarations

All authors confirm that we have complied with all relevant ethical regulations. This project was approved by the Ethics Committee of Universiti Malaysia Sabah (UMS) [JKEtika 3/23(13)] and the Animal Ethics Committee of UMS (AEC 007/2023).

## Acknowledgements

The work was partly supported by the Ministry of Higher Education Fundamental Research Grant Scheme (FRGS) (FRGS/1/2021/STG03/KDUPG/02/1).

## Author Contributions

SQO conceived and designed research. SQO, IHI and RP conducted experiments. SQO and GN, analyzed data. SQO wrote the initial draft manuscript. All authors read and approved the final manuscript.

